# AAV mediated delivery of a novel anti-BACE1 VHH reduces Abeta in an Alzheimer’s disease mouse model

**DOI:** 10.1101/698506

**Authors:** Melvin Y. Rincon, Lujia Zhou, Catherine Marneffe, Iryna Voytyuk, Yessica Wouters, Maarten Dewilde, Sandra I. Duqué, Cécile Vincke, Yona Levites, Todd E. Golde, Serge Muyldermans, Bart De Strooper, Matthew G. Holt

## Abstract

Single domain antibodies (VHH) are potentially disruptive therapeutics, with important biological value for treatment of several diseases, including neurological disorders. However, VHH have not been widely used in the central nervous system (CNS), as it is hard to reach therapeutic levels, both because of their restricted blood-brain-barrier penetration and their apparent rapid clearance from the parenchyma. Here, we propose a gene transfer strategy based on adeno-associated virus (AAV)-based vectors to deliver VHH directly into the CNS, ensuring continuous production at therapeutic levels. As a proof-of-concept, we explored the potential of AAV-delivered VHH to inhibit BACE1, a well-characterized target in Alzheimer’s disease. First, we generated a panel of VHHs targeting BACE1. One of them, VHH-B9, showed high selectivity for BACE1 and efficacy in lowering BACE1 activity in vitro. We then went on to demonstrate significant reductions in amyloid beta (Aβ) levels after AAV-based delivery of VHH-B9 into the CNS of a mouse model of cerebral amyloidosis. These results constitute a novel therapeutic approach for neurodegenerative diseases, which is applicable to a range of CNS disease targets.

## INTRODUCTION

The high selectivity of monoclonal antibodies (mAbs) offers unique opportunities to target key proteins involved in the etiology of neurodegenerative conditions, such as Parkinson’s disease and Alzheimer’s disease (AD) (Zhou *et al*, 2011; Panza *et al*, 2014). However, their potential as central nervous system (CNS) therapeutics is largely limited by their inability to cross the blood brain barrier (BBB) (Zafir-Lavie *et al*, 2018), comparatively poor biodistribution through the parenchyma (Freskgård & Urich, 2017), and short half-life (Wang *et al*, 2018). In addition, there is the potential for Fc receptor-mediated immunogenicity, mediated by microglia, which can cause vasogenic edema and cerebral microhemorrhage (Panza *et al*, 2014).

Single variable domain antibodies (VHHs) are increasingly seen as an alternative to mAbs for therapeutic use (Steeland *et al*, 2016; Bannas *et al*, 2017; Gomes *et al*, 2018). In fact, the VHH-based therapeutic Cablivi^®^ (caplacizumab-yhdp), was recently approved for market by the US Food and Drug Administration (Scully et al, 2019) for peripheral treatment of adults with acquired Thrombotic Thrombocytopenic Purpura (aTTP)

One key reason for their attractiveness over conventional mAbs is their small antigen binding site, which allows them to access unique epitopes not available to conventional mAbs, such as enzyme active sites, which are often key drug targets (Hassanzadeh-Ghassabeh *et al*, 2013). In addition, VHH have a much lower immunogenic profile than traditional mAbs, largely due to the lack of an Fc-region.

Although there have been reports that VHH can pass through the BBB (Li *et al*, 2016), the degree of penetration is variable and linked to the intrinsic charge on the protein (Bélanger *et al*, 2019). Unfortunately, the amount reaching the CNS, following peripheral injection, is further compromised by their high rate of peripheral clearance, due to renal excretion (Gainkam *et al*, 2008; Bannas *et al*, 2015). Although VHHs which do actually reach the CNS (or are directly injected into the CNS) typically have a longer half-life than in plasma, detectable levels are still reduced by up to 50% twenty-four hours post-injection (Dorresteijn *et al*, 2015). Genetic delivery of therapeutic VHHs, directly to the CNS, offers a potential solution to these issues, allowing long-term local production (Zafir-Lavie *et al*, 2018).

Adeno-associated virus (AAV)-based vectors are becoming the vehicle of choice for gene therapy applications, due to their high efficiency of gene transfer and excellent safety profile (reviewed extensively in Hudry & Vandenberghe, 2019). The major drawback of AAV vectors is the limited cargo capacity: transgenes larger than 5kb in size are not efficiently packed into the vector (Trapani *et al*, 2014). Therefore, VHH are ideal candidates for AAV-mediated delivery, as they are comparatively small (typically 350 bp in size) and can be easily incorporated into AAV vectors without extensive modifications that can adversely affect their binding properties (Pain *et al*, 2015), which is a common issue with AAV-mediated delivery of mAbs (Wu *et al*, 2010). In theory, it also allows for additional engineering, for example, directing the VHH into specific trafficking pathways to improve target engagement (Dmitriev *et al*, 2016), or incorporating specific proteolysis-promoting sequences to stimulate intracellular degradation of toxic species (Baudisch *et al*, 2018).

In this work, we describe an efficient AAV-based system for delivery of therapeutic VHH into the CNS. As a proof-of-concept strategy, we targeted the β-site amyloid precursor protein-cleaving enzyme 1 (BACE1), which is a key component in amyloid beta peptide (Aβ) production in AD (Vassar *et al*, 1999; Cai *et al*, 2001). In a two-stage process, we first generated a novel VHH, named B9, which effectively inhibits BACE1 *in vitro*. In the second phase, we then used an AAV vector to deliver B9 into the CNS of an AD mouse model, producing a long-term reduction in Aβ production after a single injection. Together, our results demonstrate that VHH against key CNS disease targets can be produced and delivered efficiently using gene therapy vectors. This approach constitutes a unique therapeutic avenue, not only for AD but for the broad spectrum of CNS diseases.

## RESULTS

### Production and identification of anti-BACE1 VHH

To generate VHH, a dromedary and a llama were immunized with purified human BACE1 ectodomain (amino acids 46-460) (Figure 1A). In total, 16 different clones were identified that bind to BACE1.

**Figure 1.**
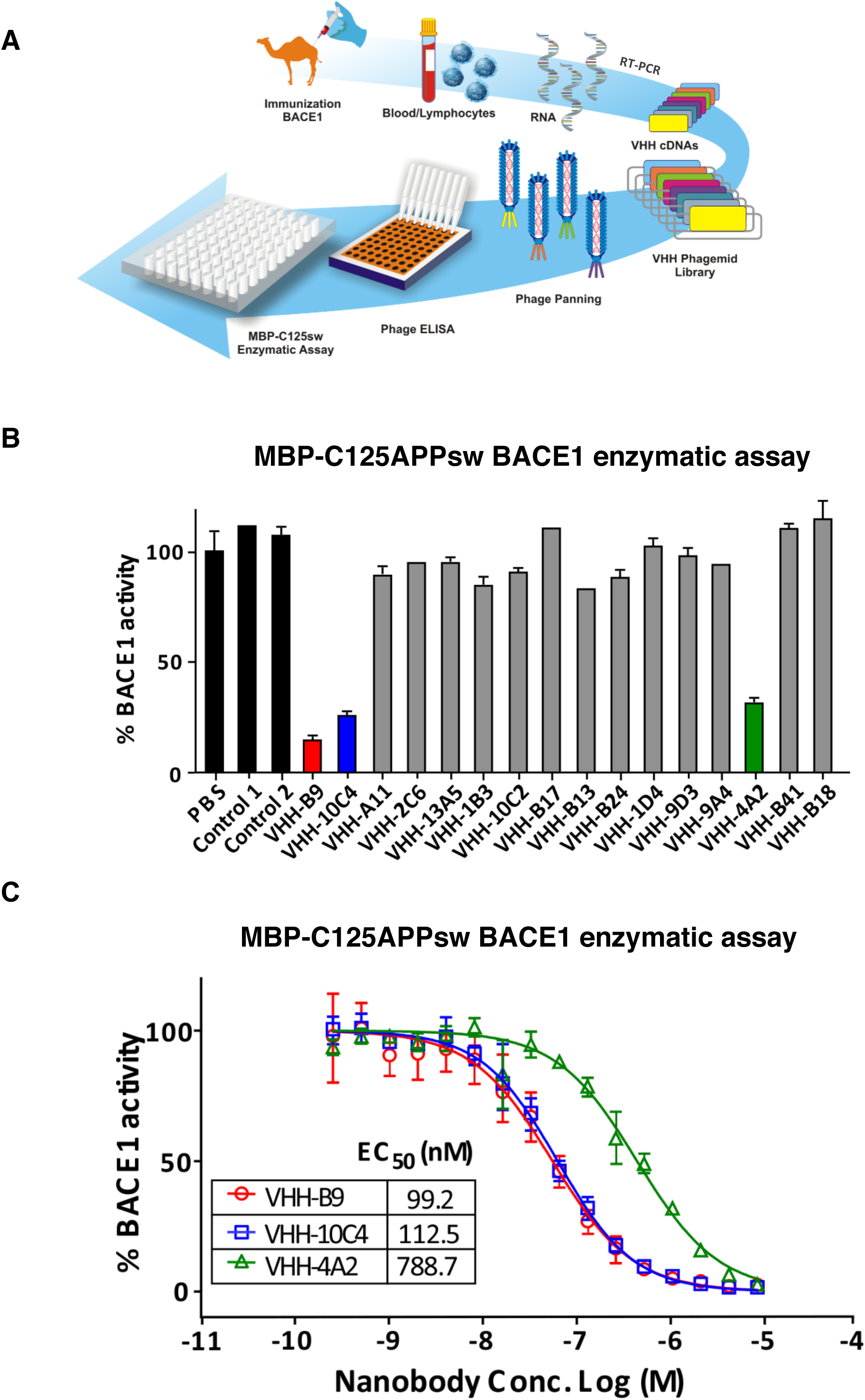
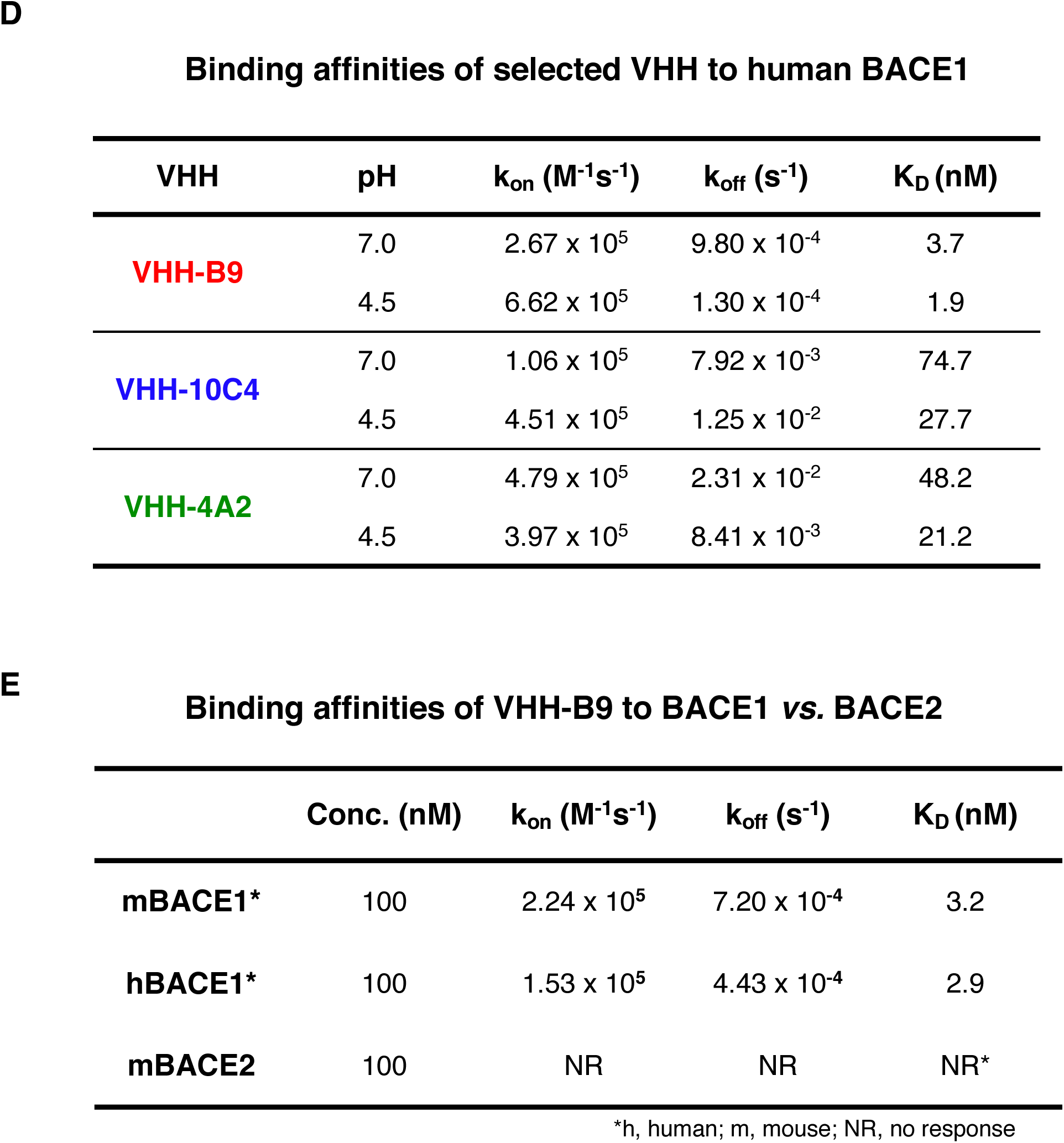
Generation and *in vitro* characterization of anti-BACE1 VHH. A. Schematic summarizing the procedure used to generate anti-BACE1 VHHs. A dromedary and a llama were immunized with recombinant human BACE1 ectodomain (amino acids 46-460). Blood lymphocytes from the immunized animals were collected for RNA extraction. cDNA was prepared and the variable fragments of heavy chain only IgGs were amplified by RT-PCR and purified using agarose gel electrophoresis. cDNAs encoding VHH were cloned into the pHEN4 phagemid vector and phage libraries were prepared. Three rounds of consecutive phage panning were performed to enrich phage particles that bound recombinant BACE1 in an ELISA assay. Binding to BACE1 was confirmed in a phage ELISA. Finally, BACE1 binding VHHs were purified and further characterized by the MBP-C125sw enzymatic assay. B. VHHs inhibiting BACE1 were identified with an *in vitro* assay, which uses the maltose binding protein (MBP) fused to a fragment of human APP containing the *Swedish* mutation (APP amino acids 571-965: K670M/N671L) (MBP-C125APPsw assay). VHHs were recombinantly expressed in bacteria, purified and added to the assay at a final concentration of 5 µM. VHH-B9, VHH-10C4 and VHH-4A2 consistently inhibited BACE1 activity, compared to PBS or control VHH (Aβ3 and BCIILP, raised against Aβ peptide and β-lactamase BCII 659/H, respectively). Values are mean ± range, n=2. C. Dose-response curves for anti-BACE1 VHHs in the MBP-C125APPsw enzymatic assay. The EC_50_ values (95% confidence interval) for VHH-B9, VHH-10C4 and VHH-4A2 are 99.2 nM (83.8-117.5 nM), 112.5 nM (101.1-125.2 nM) and 788.7 nM (682.7-911.2 nM), respectively. Values are mean ± S.D., n=3. D. Binding affinities of selected VHHs for BACE1. Measurements were made on a BIAcore instrument, using purified BACE1 ectodomain coupled to a CM5 chip. Binding constants were calculated at both pH 7.0 and pH 4.5 to confirm that the selected VHHs bind BACE1 under physiological conditions (extracellular space or trafficking endosome, respectively). E. Binding affinities of VHH-B9 for BACE1 and BACE2.

### Identification of VHH inhibiting BACE1

We screened for a possible effect of VHH on enzyme activity employing an *in vitro* APP cleavage assay (Zhou *et al*, 2011). Sixteen VHHs were recombinantly expressed in bacteria, purified and added to the assay at a final concentration of 5 µM. VHH inhibited BACE1 to varying degrees, with three VHH consistently giving the highest level of inhibition, with half-maximal effective concentrations (EC_50_) in the nanomolar range: 99.2 nM (VHH-B9), 112.5 nM (VHH-10C4) and 788.7 nM (VHH-4A2) (Figure 1B, C; Appendix Figure S1).

### Characterization of VHH-BACE1 binding

More detailed examination of VHH-BACE1 binding was obtained using the surface plasmon resonance (SPR) technique. The equilibrium dissociation constant (K_D_) was determined for VHH-B9, VHH-10C4 and VHH-4A2 at both pH 7.0 (representing the pH of the extracellular environment) and pH 4.5 (pH of the endosomal compartment). In both cases, VHH-B9 showed the highest affinity to BACE1 (Figure 1D). Interestingly, binding affinity seemed to be slightly stronger at pH 4.5, which is beneficial, as the majority of APP cleavage is reported to occur in the endosomal system (Sannerud *et al*, 2011; Ben Halima *et al*, 2016). Despite the high binding affinity of VHH-B9 towards human and mouse BACE1, it did not show any binding to mouse BACE2 under the same measurement conditions (Figure 1E), indicating a high degree of selectivity. Based on these results, VHH-B9 was chosen as the principal candidate for further characterization and testing.

We also explored the epitope on BACE1 mediating VHH-B9 binding by performing competition assays with the well-characterized monoclonal antibody 1A11 (Zhou *et al*, 2011). 1A11 was able to compete for BACE1 binding with VHH-B9 suggesting that these two antibodies bind to overlapping epitopes (Appendix Figure S2). The identity of the epitope was established by observing the binding of VHH-B9 to a series of BACE1 mutants in a dot blot assay. Our results suggest that VHH-B9 binds to BACE1 at Helix A and Loop F, which are close to the active site of the enzyme (Appendix Figure S3). Thus, VHH-B9 may act by blocking effective substrate entry

### VHH-B9 effectively inhibits neuronal BACE1 in primary neuronal cultures

To test if VHH-B9 inhibits the enzyme in its native membrane environment, we turned to primary neuronal cultures. Cells were transduced by Semliki Forest Virus (SFV) expressing wild type human APP_695_ and then treated for 12 hours with recombinant VHH-B9 at a final concentration of 3 µM. Treatment with VHH-B9 decreased the levels of sAPP_β_, CTF_β_, Aβ_1-40_ and Aβ_1-42_ detected. The levels of full-length APP, however, remained unchanged (Figure 2A, B, C). VHH-B9 inhibits BACE1 in a dose dependent manner (Figure 2D), with an EC_50_ (95% confidence interval) of 85.4 nM (52.5 nM to 125.6 nM) (Figure 2E). Together, these results indicate that VHH-B9 inhibits BACE1 activity in a native neuronal environment. Note that VHH-10C4 and VHH-4A2 were also tested in neuronal cultures. While both these VHHs inhibited BACE1 activity, VHH-B9 was the most effective, as predicted from the *in vitro* assays (Figure 1).

**Figure 2.**
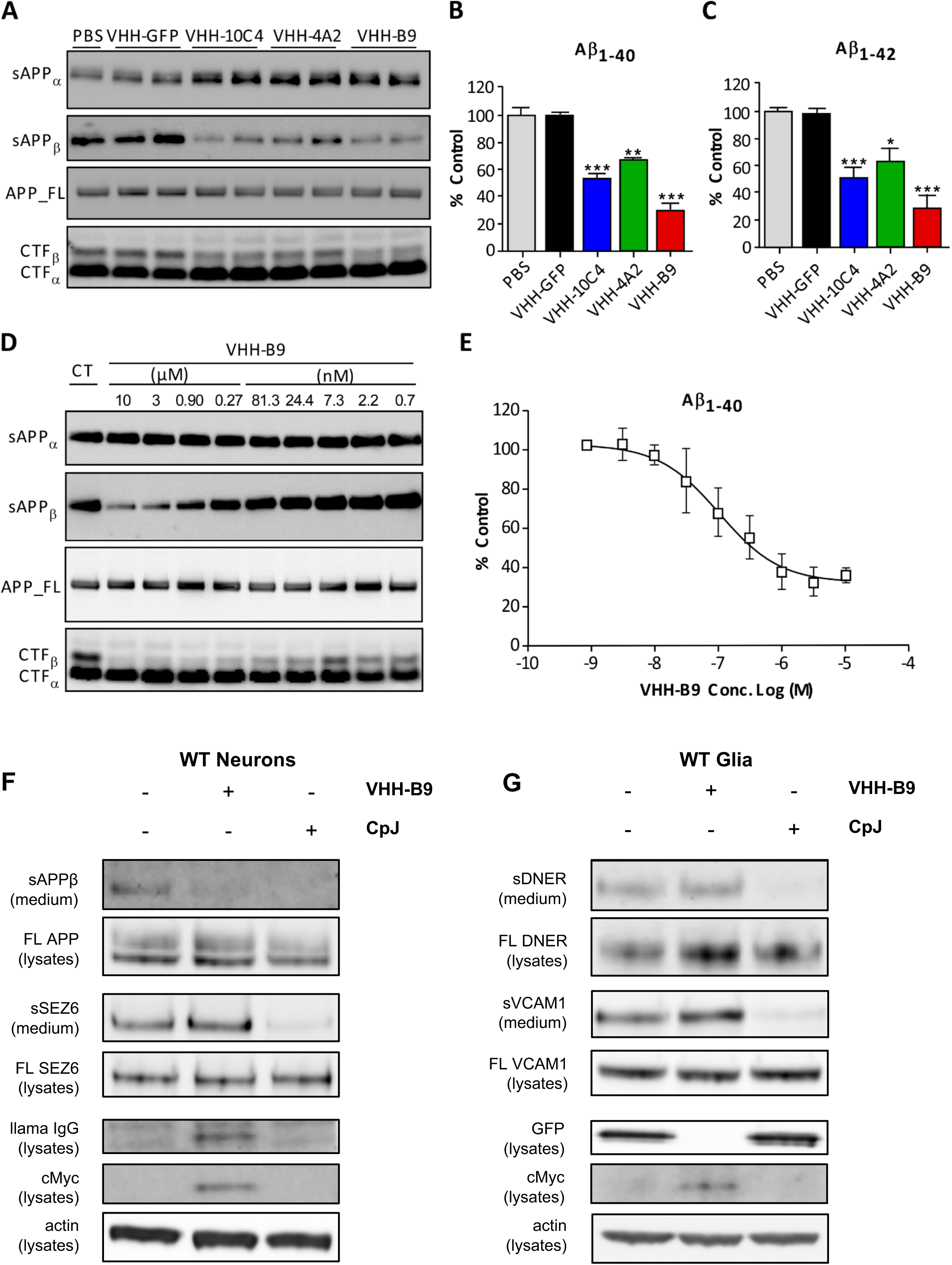
Specific inhibition of BACE1-mediated APP cleavage in primary neuronal cultures using VHH. A, B, C. Primary cultured neurons were transduced by Semliki Forest virus (SFV) expressing wild type human APP695 and treated with 3 µM of the indicated VHHs. PBS and anti-GFP VHH were used as controls. (A) Western blot analysis of conditioned media for sAPPα and sAPPβ, as well as cell extracts for full length APP, CTFα and CTFβ. (B) ELISA measurement of Aβ_1-40_ in conditioned media. (C) ELISA measurement of Aβ_1-42_ in conditioned media. Values are mean ± S.E.M., n=3 cultures for each analysis. One-way ANOVA, *** p<0.0001, **p<0.01, *p <0.5. D, E. Dose-dependent inhibition of BACE1 in primary cultured neurons by VHH-B9. Cultured neurons were transduced by SFV expressing wild type human APP695 and treated with PBS (control: CT) or decreasing concentrations of VHH-B9, ranging from 10 µM to 0.7 nM. (D) Western blot analysis of conditioned media for sAPPα and sAPPβ, as well as cell extracts for full length APP, CTFα and CTFβ. (E) Conditioned media was analyzed by ELISA to assess levels of Aβ_1-40_. Values are mean ± S.D., n=3 cultures for each analysis. The EC_50_ value (95 % confidence interval) was estimated as 85.4 nM (52.5 - 125.6 nM). F. Primary cultured neurons were transduced with an AAV vector driving the expression of VHH-B9. Compound J (CpJ) was used as control. VHH-B9 inhibited APP cleavage, as seen by a decrease in sAPPβ production. However, no decrease in SEZ6 shedding was observed. In contrast CpJ successfully inhibited both APP and SEZ6 shedding. Actin was used as a loading control. Anti-cMyc tag and anti-llama IgG were used for VHH detection. n=3 cultures for each analysis. G. Primary cultured glia were transduced with an AAV vector driving the expression of VHH-B9, or treated with CpJ. VHH-B9 had no effect on the cleavage of the BACE2 substrates DNER and VCAM1. CpJ effectively blocked shedding of both substrates. n=3 cultures for each analysis.

We next investigated the relative levels of BACE1-mediated cleavage of known substrates following treatment with VHH-B9 (Figure 2F, G). For these experiments, the cDNA for VHH-B9 was modified to contain an N-terminal BACE1 signal peptide and a C-terminal cMyc tag. This cDNA sequence was then cloned into a single-stranded AAV expression cassette and packaged into an AAV capsid (AAV-VHH-B9) (Fripont *et al*, 2019), which was used to transduce primary neuronal cultures. The non-selective BACE1 and BACE2 inhibitor Compound J (CpJ) (Esterházy *et al*, 2011) was used as a positive control. Blocking of sAPPβ shedding was observed in neurons upon transduction with AAV-VHH-B9 or CpJ treatment. In contrast, shedding of seizure protein 6 (SEZ6), a well-known neuronal substrate of BACE1 (Pigoni *et al*, 2016), was only inhibited by CpJ and remained unaffected by VHH-B9 (Figure 1F). Next, using primary glial cultures we evaluated the effects of VHH-B9 on BACE2 by assessing the cleavage of two known substrates, Delta and Notch-like epidermal growth factor-related receptor (DNER) and Vascular cell adhesion molecule 1 (VCAM1) (Voytyuk *et al*, 2018b). As expected from our previous binding data (Figure 1E), the shedding of both DNER and VCAM1 was unaffected after transduction with AAV-VHH-B9, whereas CpJ efficiently blocked the shedding of both substrates (Figure 2G).

### Viral vector mediated delivery of VHH-B9 reduced Aβ levels in a mouse model of amyloidosis

We next attempted to reduce Aβ production *in vivo* using AAV-based delivery of VHH-B9. In these experiments, an AAV vector encoding GFP was produced for use as a control. AAV vectors were tested in the APPDutch mouse model of cerebral amyloidosis. This is a transgenic line with neuronal overexpression of human E693Q APP, which causes hereditary cerebral hemorrhage with Dutch type amyloidosis. APPDutch mice present an increased Aβ_1-40/42_ ratio from an early age (Herzig *et al.*, 2004). Hence, we evaluated the activity of VHH-B9 by direct delivery of AAV-VHH-B9 (n=10) or AAV-GFP (control, n=11) vectors in six-weeks-old mice using a bilateral injection of 2 × 10^10^ vector genomes (vg) per injection site (Appendix Figure S4). Three weeks post-delivery, mice were euthanized. In animals showing VHH-B9 expression, colocalization of VHH-B9 with BACE1 was observed (Figure 3A-D), suggesting significant target engagement. ELISA measurements showed that VHH-B9 expression led to a significant decrease in levels of both Aβ_1-40_ and Aβ_1-42_ (Figure 3E, F), indicative of BACE1 inhibition. No apparent toxicity, derived from either the VHH or the vector, was observed, either on animal survival rates, or at the histological level post-mortem (data not shown).

**Figure 3.**
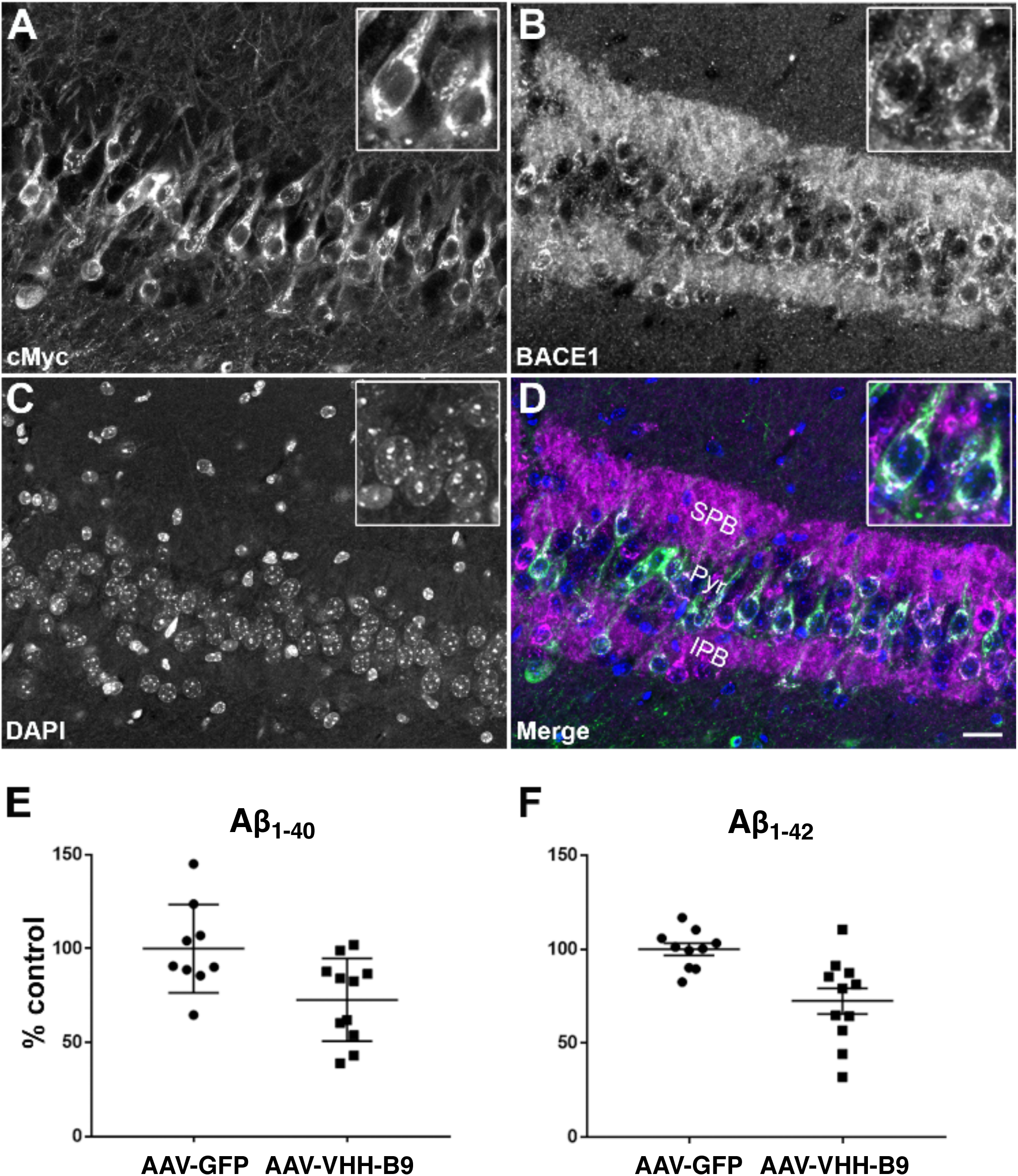
Long-term BACE1 inhibition in APPDutch mice after AAV-VHH delivery significantly reduces amyloid beta load. A-D. AAV-VHH-B9 vector was injected into the hippocampus of APPDutch mice (n=3) at a dose of 2×10^10^ vg per site. Three weeks post-injection, brains were recovered and processed for immunohistochemistry. Representative images of coronal brain sections are shown for staining against BACE1 (magenta) and cMyc (green). Images show colocalization of VHH-B9 with BACE1 in the pyramidal cell layer (Pyr), as well as in the suprapyramidal blade (SPB) and infrapyramidal blade (IPB) of the mossy fibers. The diffuse staining patterns suggest that VHH-B9 is engaged with BACE1 in internal structures, such as early endosomes. Scale bar, 25 µm. E, F. APP Dutch mice were injected in the hippocampus with AAV-VHH-B9 (n=11) or the control vector AAV-GFP (n=10). Whole brains were collected 3 weeks post-injection for analysis. Soluble protein was extracted and used for quantification of Aβ_1-40_ and Aβ_1-42_ levels by ELISA. VHH-B9 expression led to a significant decrease in both Aβ_1-40_ (27.3%, p=0.0168) and Aβ_1-42_ (27.5%, p=0.0026) levels.

## DISCUSSION

We exploited the known strengths of the VHH platform to successfully demonstrate targeting of BACE1, an important CNS target central to AD pathology (Vassar *et al*, 1999; Barão *et al*, 2016; Voytyuk *et al*, 2018b). These strengths include the ability to rapidly and easily identify VHHs that specifically bind the intended target with adequate affinity, under physiological conditions (Steeland *et al*, 2016). Furthermore, by exploiting the high intrinsic stability of VHHs and ease of engineering (Vincke *et al*, 2009), we were able to incorporate a signal peptide into the VHH sequence, allowing intracellular expression and directed trafficking to the endosome, where the majority of BACE1-mediated APP cleavage is thought to occur (Sannerud *et al*, 2011; Ben Halima *et al*, 2016). This was done with the aim of promoting VHH-BACE1 interactions early in the secretory pathway, which may be beneficial as some mutations in APP, such as the *Swedish* mutation (*Swe*), are known to promote processing of APP in the secretory pathway itself (Sasaguri *et al*, 2017). Finally, the small size of VHH allowed easy incorporation into an AAV-based vector (Verhelle *et al*, 2017). Use of such a system facilitated intracellular delivery and continuous VHH expression, with direct parenchymal injection circumventing the shielding effect of the BBB, which has often limited the therapeutic efficiency of antibody therapies for brain disorders (Barão *et al*, 2016). This allowed us to achieve long-term reductions in BACE1 activity and Aβ production in a mouse model of Alzheimer’s type amyloidosis.

As BACE1 contributes to 80% of Aβ_1-40_ production in the brain (Atwal *et al*, 2011). The decreases in steady state Aβ levels that we report here are significant, especially considering they were achieved after a single administration of vector. By way of comparison, a single intracisternal injection of 75 µg VHH-B3a in APPswe/PS1dE9 mice (Dorresteijn *et al*, 2015) showed no significant decrease in the level of soluble Aβ_1-40_ twenty-four hours post-injection. This presumably reflects the relative high dissociation constant of this VHH for BACE1 (0.3 µM), a gradual decrease in available VHH (possibly as a result of clearance), or a combination of both factors. In contrast, direct injections of conventional mAbs, or systemic administration of mAbs engineered to cross the BBB by receptor-mediated transcytosis, led to BACE1 inhibition in a dose-dependent fashion (Zhou *et al*, 2011; Cheng *et al*, 2013; Ye *et al*, 2017). In these examples, however, effective inhibition was typically limited to a period of up to forty-eight hours post-injection. Taken together, we believe these results largely validate our experimental strategy, although further work will be needed to determine the optimal temporal window for intervention in the disease process.

At present, the levels of Aβ decrease necessary to produce an effect in AD are undetermined (Kennedy *et al*, 2016; Timmers *et al*, 2018; Egan *et al*, 2019). However, there are reasons to believe that a sustained steady state reduction, similar to that which we report, may be worthwhile pursuing. First, a reduction of 50% in the levels of BACE1 activity is sufficient to significantly reduce Aβ plaques, neuritic burden, and synaptic deficits in mouse models of AD (McConlogue *et al*, 2007), similar to the effects seen with a reduction of 30% in the levels of gamma secretase activity (Li *et al*, 2007). Second, a mutation in human APP, which reduces cleavage by BACE1 (Jonsson *et al*, 2012), leads to a reduction in cerebral Aβ of approximately 20%, with carriers showing lifelong protection against AD and cognitive decline (although issues with the mutation affecting aggregation of Aβ peptides cannot be fully excluded) (Maloney *et al*, 2014; Benilova *et al*, 2014).

Sustained protection through low to moderate levels of CNS-localized BACE1-specific inhibition may well have the additional benefit of reducing, or avoiding, toxicity related to complete loss of BACE1 function, such as hypomyelination (Willem *et al*, 2006), aberrant synaptic homeostasis and plasticity (Filser *et al*, 2015), axon guidance abnormalities (Rajapaksha *et al*, 2011; Cao *et al*, 2012; Ou-Yang *et al*, 2018), impairments in spatial and working memory (Henley *et al*, 2019; Knopman, 2019; Egan *et al*, 2019) and retinal pathology (Cai *et al*, 2012). In this respect, it is important to note that the levels of BACE1 inhibition achieved with AAV-mediated delivery of VHH-B9 did not substantially affect cleavage of the alternative BACE1 substrate seizure protein 6 (SEZ6), which is important for maintenance of dendritic spines and synaptic plasticity (Gunnersen *et al*, 2007; Pigoni *et al*, 2016; Zhu *et al*, 2018). This is most simply explained by the relative amounts of APP and SEZ6 available for cleavage in a mass-action model (Wilhelm *et al*, 2014). Peripheral effects of BACE1 inhibition on muscle spindle assembly, as well as cross-reactivity with BACE2, Cathepsin D and Cathepsin E, which can lead to impairments in glucose homeostasis, hypopigmentation, seizures and blindness (amongst others), are also avoided (Voytyuk *et al*, 2018b). Crucially, no signs of apparent toxicity from the VHH or viral vector (at the doses used) were observed over 4 weeks post-vector delivery, consistent with predictions on tolerability from other studies (LeWitt *et al*, 2011; Saraiva *et al*, 2016). Long-term studies focusing on these issues will, however, be essential to determine the clinical feasibility of this approach.

Additional experiments will also be needed to determine the effective therapeutic dose of AAV-VHH-B9 required and whether any reduction in total Aβ load leads to improvements in cognitive function. However, if required, modifications to the system can easily be made by taking advantage of the relative ease with which VHH and AAV can be engineered. For example, VHH affinities in the low picomolar range can be achieved using standard affinity maturation techniques (Mahajan *et al*, 2018), or via production of bivalent VHH constructs (Beirnaert *et al*, 2017) that bind distinct epitopes on a given target. Expression levels of a given VHH from a vector-based system can be improved by use of a self-complementary genome configuration, containing multiple copies of the VHH coding sequence (Verhelle *et al*, 2017). Use of specific promoter systems, in combination with cell type specific transcriptional enhancers and inducible elements (Hudry & Vandenberghe, 2019), should also allow temporally controllable and graded levels of VHH expression in cell types of interest. As demonstrated, VHH can be modified to add unique functions, such as signal peptide sequences (Vincke *et al*, 2009). This can be further exploited to include, for example, sequences targeting the VHH-antigen complex for degradation (Caussinus *et al*, 2011). The efficacy of such modifications, for example in reducing Aβ levels, remains to be established. Finally, AAV vectors which cross the BBB at high efficiency and achieve widespread CNS transduction are being developed. These AAV vectors show considerable promise for delivery of therapeutics to the CNS following systemic injection, removing the need for invasive direct intraparenchymal injections or cerebrospinal fluid-based injections, whilst facilitating broad transgene delivery (Deverman *et al*, 2016). In reality, it is likely to be a combination of technological developments in the aforementioned areas that finally makes AAV-mediated delivery of antibodies, or antibody fragments, a clinically relevant option for CNS disorders.

As production costs drop, such single use AAV technology is likely to offer a more cost-effective alternative to regular infusions of recombinant antibodies or anti-sense oligonucleotides.

In summary, using the well characterized read out of BACE1inhibition as a *proof-of-concept*, we have shown the viability of combining VHH with viral vector-mediated delivery to target a specific CNS protein, producing local and statistically significant reduction in Aβ levels, after a single injection. Moving forward, we propose that vector-mediated delivery of VHHs can be successfully exploited for the treatment of a variety of CNS conditions with defined targets, such as Parkinson’s disease and amyotrophic lateral sclerosis, which are, at present, untreatable.

## ACKNOWLEDGEMENTS

Dr. Matthias Jucker kindly provided the APPDutch mouse model. MYR is a postdoctoral researcher with the FWO (133722/1204517N) and acknowledges the continuous support of the Fundación Cardiovascular de Colombia. SID was supported by The Foundation for Alzheimer Research (SAO-FRA) (P#14006). MGH was supported by a VIB institutional grant and external support from the Thierry Latran Foundation (SOD-VIP), FWO (Grant 1513616N) and European Research Council (ERC) (Starting Grant 281961 – AstroFunc; Proof of Concept Grant 713755 – AD-VIP). Work in the BDS Lab is supported by the Opening the Future campaign of the KU Leuven, SAO-FRA (P#16017), FWO, KU Leuven, VIB, a Methusalem grant from KU Leuven and the Flemish Government, the Flanders Network for Dementia Research (VIND, Strategic Basic Research Grant 135043) and the Alzheimer’s Association. BDS is supported by the Geneeskundige Stichting Koningin Elisabeth and the Bax-Vanluffelen Chair for Alzheimer’s disease. We want to thank the collaborators of the VIB Nanobody Core for their valuable contribution to the research presented in this paper.

## AUTHOR CONTRIBUTIONS

BDS and MGH conceived the project. LZ, CV, SM, YW, MD and BDS generated and characterized the anti-BACE1 VHH. MYR generated the AAV used in tissue culture experiments; YL and TEG generated the AAV used for *in vivo* experiments. LZ, CM and IV performed tissue culture experiments. MYR and SID performed experiments with APPDutch mice. MYR, LZ, SID, BDS and MGH analyzed the data. MGH, MYR and LZ wrote the manuscript with input from all authors.

## REFERENCES

Atwal JK, Chen Y, Chiu C, Mortensen DL, Meilandt WJ, Liu Y, Heise CE, Hoyte K, Luk W, Lu Y, Peng K, Wu P, Rouge L, Zhang Y, Lazarus RA, Scearce-Levie K, Wang W, Wu Y, Tessier-Lavigne M & Watts RJ (2011) A therapeutic antibody targeting BACE1 inhibits amyloid-β production in vivo. Sci Transl Med 3:84ra43

Bannas P, Hambach J & Koch-Nolte F (2017) Nanobodies and Nanobody-Based Human Heavy Chain Antibodies As Antitumor Therapeutics. Front Immunol 8: 1603

Bannas P, Lenz A, Kunick V, Fumey W, Rissiek B, Schmid J, Haag F, Leingärtner A, Trepel M, Adam G & Koch-Nolte F (2015) Validation of nanobody and antibody based in vivo tumor xenograft NIRF-imaging experiments in mice using ex vivo flow cytometry and microscopy. J Vis Exp: e52462

Barão S, Moechars D, Lichtenthaler SF & De Strooper B (2016) BACE1 Physiological Functions May Limit Its Use as Therapeutic Target for Alzheimer’s Disease. Trends Neurosci. 39: 158–169

Baudisch B, Pfort I, Sorge E & Conrad U (2018) Nanobody-Directed Specific Degradation of Proteins by the 26S-Proteasome in Plants. Front Plant Sci 9: 130

Beirnaert E, Desmyter A, Spinelli S, Lauwereys M, Aarden L, Dreier T, Loris R, Silence K, Pollet C, Cambillau C & de Haard H (2017) Bivalent Llama Single-Domain Antibody Fragments against Tumor Necrosis Factor Have Picomolar Potencies due to Intramolecular Interactions. Front Immunol 8: 867

Bélanger K, Iqbal U, Tanha J, MacKenzie R, Moreno M & Stanimirovic D (2019) Single-Domain Antibodies as Therapeutic and Imaging Agents for the Treatment of CNS Diseases. Antibodies 8: 27

Ben Halima S, Mishra S, Raja KMP, Willem M, Baici A, Simons K, Brüstle O, Koch P, Haass C, Caflisch A & Rajendran L (2016) Specific Inhibition of β-Secretase Processing of the Alzheimer Disease Amyloid Precursor Protein. Cell Rep 14: 2127–2141

Benilova I, Gallardo R, Ungureanu A-A, Castillo Cano V, Snellinx A, Ramakers M, Bartic C, Rousseau F, Schymkowitz J & De Strooper B (2014) The Alzheimer disease protective mutation A2T modulates kinetic and thermodynamic properties of amyloid-β (Aβ) aggregation. J. Biol. Chem. 289: 30977–30989

Cai H, Wang Y, McCarthy D, Wen H, Borchelt DR, Price DL & Wong PC (2001) BACE1 is the major beta-secretase for generation of Abeta peptides by neurons. Nat. Neurosci. 4: 233–234

Cai J, Qi X, Kociok N, Skosyrski S, Emilio A, Ruan Q, Han S, Liu L, Chen Z, Bowes Rickman C, Golde T, Grant MB, Saftig P, Serneels L, de Strooper B, Joussen AM & Boulton ME (2012) β-Secretase (BACE1) inhibition causes retinal pathology by vascular dysregulation and accumulation of age pigment. EMBO Mol Med 4: 980–991

Cao L, Rickenbacher GT, Rodriguez S, Moulia TW & Albers MW (2012) The precision of axon targeting of mouse olfactory sensory neurons requires the BACE1 protease. Sci Rep 2: 231

Caussinus E, Kanca O & Affolter M (2011) Fluorescent fusion protein knockout mediated by anti-GFP nanobody. Nat. Struct. Mol. Biol. 19: 117–121

Cheng X, Zhou Y, Gu W, Wu J, Nie A, Cheng J, Zhou J, Zhou W & Zhang Y (2013) The selective BACE1 inhibitor VIa reduces amyloid-β production in cell and mouse models of Alzheimer’s disease. J. Alzheimers Dis. 37: 823–834

Deverman BE, Pravdo PL, Simpson BP, Kumar SR, Chan KY, Banerjee A, Wu W-L, Yang B, Huber N, Pasca SP & Gradinaru V (2016) Cre-dependent selection yields AAV variants for widespread gene transfer to the adult brain. Nat. Biotechnol. 34: 204–209

Dmitriev OY, Lutsenko S & Muyldermans S (2016) Nanobodies as Probes for Protein Dynamics in Vitro and in Cells. J. Biol. Chem. 291: 3767–3775

Dorresteijn B, Rotman M, Faber D, Schravesande R, Suidgeest E, van der Weerd L, van der Maarel SM, Verrips CT & El Khattabi M (2015) Camelid heavy chain only antibody fragment domain against β-site of amyloid precursor protein cleaving enzyme 1 inhibits β-secretase activity in vitro and in vivo. FEBS J. 282: 3618–3631

Egan MF, Kost J, Voss T, Mukai Y, Aisen PS, Cummings JL, Tariot PN, Vellas B, van Dyck CH, Boada M, Zhang Y, Li W, Furtek C, Mahoney E, Harper Mozley L, Mo Y, Sur C & Michelson D (2019) Randomized Trial of Verubecestat for Prodromal Alzheimer’s Disease. N. Engl. J. Med. 380: 1408–1420

Esterházy D, Stützer I, Wang H, Rechsteiner MP, Beauchamp J, Döbeli H, Hilpert H, Matile H, Prummer M, Schmidt A, Lieske N, Boehm B, Marselli L, Bosco D, Kerr-Conte J, Aebersold R, Spinas GA, Moch H, Migliorini C & Stoffel M (2011) Bace2 Is a β Cell-Enriched Protease that Regulates Pancreatic β Cell Function and Mass. Cell Metabolism 14: 365–377

Filser S, Ovsepian SV, Masana M, Blazquez-Llorca L, Brandt Elvang A, Volbracht C, Müller MB, Jung CKE & Herms J (2015) Pharmacological inhibition of BACE1 impairs synaptic plasticity and cognitive functions. Biol. Psychiatry 77: 729–739

Freskgård P-O & Urich E (2017) Antibody therapies in CNS diseases. Neuropharmacology 120: 38–55

Fripont S, Marneffe C, Marino M, Rincon MY & Holt MG (2019) Production, Purification, and Quality Control for Adeno-associated Virus-based Vectors. J Vis Exp

Gainkam LOT, Huang L, Caveliers V, Keyaerts M, Hernot S, Vaneycken I, Vanhove C, Revets H, De Baetselier P & Lahoutte T (2008) Comparison of the biodistribution and tumor targeting of two 99mTc-labeled anti-EGFR nanobodies in mice, using pinhole SPECT/micro-CT. J. Nucl. Med. 49: 788–795

Gomes JR, Cabrito I, Soares HR, Costelha S, Teixeira A, Wittelsberger A, Stortelers C, Vanlandschoot P & Saraiva MJ (2018) Delivery of an anti-transthyretin Nanobody to the brain through intranasal administration reveals transthyretin expression and secretion by motor neurons. J. Neurochem. 145: 393–408

Gunnersen JM, Kim MH, Fuller SJ, De Silva M, Britto JM, Hammond VE, Davies PJ, Petrou S, Faber ESL, Sah P & Tan S-S (2007) Sez-6 proteins affect dendritic arborization patterns and excitability of cortical pyramidal neurons. Neuron 56: 621–639

Hassanzadeh-Ghassabeh G, Devoogdt N, De Pauw P, Vincke C & Muyldermans S (2013) Nanobodies and their potential applications. Nanomedicine (Lond) 8: 1013–1026

Henley D, Raghavan N, Sperling R, Aisen P, Raman R & Romano G (2019) Preliminary Results of a Trial of Atabecestat in Preclinical Alzheimer’s Disease. N. Engl. J. Med. 380: 1483–1485

Hudry E & Vandenberghe LH (2019) Therapeutic AAV Gene Transfer to the Nervous System: A Clinical Reality. Neuron 101: 839–862

Jonsson T, Atwal JK, Steinberg S, Snaedal J, Jonsson PV, Bjornsson S, Stefansson H, Sulem P, Gudbjartsson D, Maloney J, Hoyte K, Gustafson A, Liu Y, Lu Y, Bhangale T, Graham RR, Huttenlocher J, Bjornsdottir G, Andreassen OA, Jönsson EG, et al (2012) A mutation in APP protects against Alzheimer’s disease and age-related cognitive decline. Nature 488: 96–99

Kennedy ME, Stamford AW, Chen X, Cox K, Cumming JN, Dockendorf MF, Egan M, Ereshefsky L, Hodgson RA, Hyde LA, Jhee S, Kleijn HJ, Kuvelkar R, Li W, Mattson BA, Mei H, Palcza J, Scott JD, Tanen M, Troyer MD, et al (2016) The BACE1 inhibitor verubecestat (MK-8931) reduces CNS β-amyloid in animal models and in Alzheimer’s disease patients. Sci Transl Med 8: 363ra150

Kimura R, Devi L & Ohno M (2010) Partial reduction of BACE1 improves synaptic plasticity, recent and remote memories in Alzheimer’s disease transgenic mice. J. Neurochem. 113: 248–261

Knopman DS (2019) Lowering of Amyloid-Beta by β-Secretase Inhibitors - Some Informative Failures. N. Engl. J. Med. 380: 1476–1478

LeWitt PA, Rezai AR, Leehey MA, Ojemann SG, Flaherty AW, Eskandar EN, Kostyk SK, Thomas K, Sarkar A, Siddiqui MS, Tatter SB, Schwalb JM, Poston KL, Henderson JM, Kurlan RM, Richard IH, Van Meter L, Sapan CV, During MJ, Kaplitt MG, et al (2011) AAV2-GAD gene therapy for advanced Parkinson’s disease: a double-blind, sham-surgery controlled, randomised trial. The Lancet Neurology 10: 309–319

Li T, Bourgeois J-P, Celli S, Glacial F, Le Sourd A-M, Mecheri S, Weksler B, Romero I, Couraud P-O, Rougeon F & Lafaye P (2012) Cell-penetrating anti-GFAP VHH and corresponding fluorescent fusion protein VHH-GFP spontaneously cross the blood-brain barrier and specifically recognize astrocytes: application to brain imaging. FASEB J. 26: 3969–3979

Li T, Vandesquille M, Koukouli F, Dudeffant C, Youssef I, Lenormand P, Ganneau C, Maskos U, Czech C, Grueninger F, Duyckaerts C, Dhenain M, Bay S, Delatour B & Lafaye P (2016) Camelid single-domain antibodies: A versatile tool for in vivo imaging of extracellular and intracellular brain targets. J Control Release 243: 1–10

Li T, Wen H, Brayton C, Laird FM, Ma G, Peng S, Placanica L, Wu TC, Crain BJ, Price DL, Eberhart CG & Wong PC (2007) Moderate Reduction of γ-Secretase Attenuates Amyloid Burden and Limits Mechanism-Based Liabilities. J. Neurosci. 27: 10849–10859

Mahajan SP, Meksiriporn B, Waraho-Zhmayev D, Weyant KB, Kocer I, Butler DC, Messer A, Escobedo FA & DeLisa MP (2018) Computational affinity maturation of camelid single-domain intrabodies against the nonamyloid component of alpha-synuclein. Sci Rep 8: 17611

Maloney JA, Bainbridge T, Gustafson A, Zhang S, Kyauk R, Steiner P, van der Brug M, Liu Y, Ernst JA, Watts RJ & Atwal JK (2014) Molecular mechanisms of Alzheimer disease protection by the A673T allele of amyloid precursor protein. J. Biol. Chem. 289: 30990–31000

McConlogue L, Buttini M, Anderson JP, Brigham EF, Chen KS, Freedman SB, Games D, Johnson-Wood K, Lee M, Zeller M, Liu W, Motter R & Sinha S (2007) Partial reduction of BACE1 has dramatic effects on Alzheimer plaque and synaptic pathology in APP Transgenic Mice. J. Biol. Chem. 282: 26326–26334

Ou-Yang M-H, Kurz JE, Nomura T, Popovic J, Rajapaksha TW, Dong H, Contractor A, Chetkovich DM, Tourtellotte WG & Vassar R (2018) Axonal organization defects in the hippocampus of adult conditional BACE1 knockout mice. Sci Transl Med 10: 459

Pain C, Dumont J & Dumoulin M (2015) Camelid single-domain antibody fragments: Uses and prospects to investigate protein misfolding and aggregation, and to treat diseases associated with these phenomena. Biochimie 111: 82–106

Panza F, Solfrizzi V, Imbimbo BP & Logroscino G (2014) Amyloid-directed monoclonal antibodies for the treatment of Alzheimer’s disease: the point of no return? Expert Opin Biol Ther 14: 1465–1476

Pigoni M, Wanngren J, Kuhn P-H, Munro KM, Gunnersen JM, Takeshima H, Feederle R, Voytyuk I, De Strooper B, Levasseur MD, Hrupka BJ, Müller SA & Lichtenthaler SF (2016) Seizure protein 6 and its homolog seizure 6-like protein are physiological substrates of BACE1 in neurons. Mol Neurodegener 11: 67

Rajapaksha TW, Eimer WA, Bozza TC & Vassar R (2011) The Alzheimer’s β-secretase enzyme BACE1 is required for accurate axon guidance of olfactory sensory neurons and normal glomerulus formation in the olfactory bulb. Mol Neurodegener 6: 88

Sannerud R, Declerck I, Peric A, Raemaekers T, Menendez G, Zhou L, Veerle B, Coen K, Munck S, De Strooper B, Schiavo G & Annaert W (2011) ADP ribosylation factor 6 (ARF6) controls amyloid precursor protein (APP) processing by mediating the endosomal sorting of BACE1. Proc. Natl. Acad. Sci. U.S.A. 108: E559–568

Saraiva J, Nobre RJ & Pereira de Almeida L (2016) Gene therapy for the CNS using AAVs: The impact of systemic delivery by AAV9. J Control Release 241: 94–109

Sasaguri H, Nilsson P, Hashimoto S, Nagata K, Saito T, Strooper BD, Hardy J, Vassar R, Winblad B & Saido TC (2017) APP mouse models for Alzheimer’s disease preclinical studies. The EMBO Journal 36: 2473–2487

Steeland S, Vandenbroucke RE & Libert C (2016) Nanobodies as therapeutics: big opportunities for small antibodies. Drug Discovery Today 21: 1076–1113

Timmers M, Streffer JR, Russu A, Tominaga Y, Shimizu H, Shiraishi A, Tatikola K, Smekens P, Börjesson-Hanson A, Andreasen N, Matias-Guiu J, Baquero M, Boada M, Tesseur I, Tritsmans L, Van Nueten L & Engelborghs S (2018) Pharmacodynamics of atabecestat (JNJ-54861911), an oral BACE1 inhibitor in patients with early Alzheimer’s disease: randomized, double-blind, placebo-controlled study. Alzheimers Res Ther 10: 85

Trapani I, Colella P, Sommella A, Iodice C, Cesi G, de Simone S, Marrocco E, Rossi S, Giunti M, Palfi A, Farrar GJ, Polishchuk R & Auricchio A (2014) Effective delivery of large genes to the retina by dual AAV vectors. EMBO Mol Med 6: 194–211

Vassar R, Bennett BD, Babu-Khan S, Kahn S, Mendiaz EA, Denis P, Teplow DB, Ross S, Amarante P, Loeloff R, Luo Y, Fisher S, Fuller J, Edenson S, Lile J, Jarosinski MA, Biere AL, Curran E, Burgess T, Louis JC, et al (1999) Beta-secretase cleavage of Alzheimer’s amyloid precursor protein by the transmembrane aspartic protease BACE. Science 286: 735–741

Verhelle A, Nair N, Everaert I, Van Overbeke W, Supply L, Zwaenepoel O, Peleman C, Van Dorpe J, Lahoutte T, Devoogdt N, Derave W, Chuah MK, VandenDriessche T & Gettemans J (2017) AAV9 delivered bispecific nanobody attenuates amyloid burden in the gelsolin amyloidosis mouse model. Hum. Mol. Genet. 26: 1353–1364

Vincke C, Loris R, Saerens D, Martinez-Rodriguez S, Muyldermans S & Conrath K (2009) General strategy to humanize a camelid single-domain antibody and identification of a universal humanized nanobody scaffold. J. Biol. Chem. 284: 3273–3284

Voytyuk I, De Strooper B & Chávez-Gutiérrez L (2018a) Modulation of γ- and β-Secretases as Early Prevention Against Alzheimer’s Disease. Biol. Psychiatry 83: 320–327

Voytyuk I, Mueller SA, Herber J, Snellinx A, Moechars D, van Loo G, Lichtenthaler SF & De Strooper B (2018b) BACE2 distribution in major brain cell types and identification of novel substrates. Life Sci Alliance 1: e201800026

Wang Q, Delva L, Weinreb PH, Pepinsky RB, Graham D, Veizaj E, Cheung AE, Chen W, Nestorov I, Rohde E, Caputo R, Kuesters GM, Bohnert T & Gan L-S (2018) Monoclonal antibody exposure in rat and cynomolgus monkey cerebrospinal fluid following systemic administration. Fluids Barriers CNS 15: 10

Wang W, Liu Y & Lazarus RA (2013) Allosteric inhibition of BACE1 by an exosite-binding antibody. Current Opinion in Structural Biology 23: 797–805

Wilhelm BG, Mandad S, Truckenbrodt S, Kröhnert K, Schäfer C, Rammner B, Koo SJ, Claßen GA, Krauss M, Haucke V, Urlaub H & Rizzoli SO (2014) Composition of isolated synaptic boutons reveals the amounts of vesicle trafficking proteins. Science 344: 1023–1028

Willem M, Garratt AN, Novak B, Citron M, Kaufmann S, Rittger A, DeStrooper B, Saftig P, Birchmeier C & Haass C (2006) Control of peripheral nerve myelination by the beta-secretase BACE1. Science 314: 664–666

Wu Z, Yang H & Colosi P (2010) Effect of genome size on AAV vector packaging. Mol. Ther. 18: 80–86

Yan R & Vassar R (2014) Targeting the β secretase BACE1 for Alzheimer’s disease therapy. Lancet Neurol 13: 319–329

Ye X, Feng T, Tammineni P, Chang Q, Jeong YY, Margolis DJ, Cai H, Kusnecov A & Cai Q (2017) Regulation of Synaptic Amyloid-β Generation through BACE1 Retrograde Transport in a Mouse Model of Alzheimer’s Disease. J. Neurosci. 37: 2639–2655

Zafir-Lavie I, Sherbo S, Goltsman H, Badinter F, Yeini E, Ofek P, Miari R, Tal O, Liran A, Shatil T, Krispel S, Shapir N, Neil GA, Benhar I, Panet A & Satchi-Fainaro R (2018) Successful intracranial delivery of trastuzumab by gene-therapy for treatment of HER2-positive breast cancer brain metastases. J Control Release 291: 80–89

Zhou L, Chávez-Gutiérrez L, Bockstael K, Sannerud R, Annaert W, May PC, Karran E & De Strooper B (2011) Inhibition of beta-secretase in vivo via antibody binding to unique loops (D and F) of BACE1. J. Biol. Chem. 286: 8677–8687

Zhu K, Xiang X, Filser S, Marinković P, Dorostkar MM, Crux S, Neumann U, Shimshek DR, Rammes G, Haass C, Lichtenthaler SF, Gunnersen JM & Herms J (2018) Beta-Site Amyloid Precursor Protein Cleaving Enzyme 1 Inhibition Impairs Synaptic Plasticity via Seizure Protein 6. Biol. Psychiatry 83: 428–437

