## Supplementary Material & Methods for "AAV mediated delivery of a novel anti-BACE1 VHH reduces Abeta in an Alzheimer’s disease mouse model"

### SUPPLEMENTARY INFORMATION

#### SUPPLEMENTARY FIGURE LEGENDS

**Supplementary Figure S1. Protein sequences of anti-BACE1 VHH.**

Sequences are aligned, with framework and CDR regions indicated.

**Supplementary Figure S2. VHH-B9 and monoclonal antibody 1A11 compete for binding to BACE1.**

Epitope binning was done by biolayer interferometry (Octet RED96; Molecular Devices). Biotinylated BACE1 was bound to streptavidin sensor tips (not shown). Afterwards, BACE1 loaded sensors were dipped in 1A11, VHH-B9 or kinetic buffer for 600 seconds to allow binding. After a 30 seconds equilibration in kinetic buffer, tips were dipped in (A) VHH-B9 or (B) 1A11. (A) No additional binding of VHH-B9 was observed. (B) A low level of 1A11 binding to the BACE1-VHH-B9 complex was observed. This residual binding is most probably due to the displacement of VHH-B9 by 1A11 (which shows a stronger binding to BACE1 than VHH-B9 due to avidity-mediated mechanisms), suggesting that these two antibodies bind to the same or adjacent epitopes.

**Supplementary Figure S3. VHH-B9 binds to a unique exosite on BACE1.**

Wild type (WT) BACE1 ectodomain (1-460), or various mutants including S376Q377D378/WAA (F(SQD/WAAA)), EDVATSQDD371-379/MGAGLNYE (F(BACE2)), EVATSQD371-378/EGS (ΔF) and GFPLNQSEVLASVG219-232/GAG (ΔA), were purified from cultures of HEK293 cells. 200 ng purified protein was dotted onto a nitrocellulose membrane and probed with the indicated antibodies. Monoclonal antibodies 10B8, 5G7 and 1A11 were used as positive controls. 10B8 and 5G7 recognize all forms of BACE1 tested, indicating that the mutants were properly folded. 1A11 does not recognize BACE1 with mutations in loop F but recognizes Helix A mutants, as well as WT BACE1, as previously reported (Zhou *et al*, 2011). VHH-B9 only weakly recognized ΔLoop F and ΔHelix A mutants, but recognized other Loop F mutants, S376Q377D378/WAA and EDVATSQDD371-379/MGAGLNYE. Helix A is a structure adjacent to Loop F flanking the active-site cleft of BACE1. This suggests that VHH-B9 binds via a conformational epitope engaging Helix A and Loop F, although direct binding to Loop F may not necessarily be needed. Binding to these structural elements, unique to BACE1, likely explains the specific inhibition seen with VHH-B9.

**Supplementary Figure S4. Biodistribution of VHH-B9 in the hippocampus of wild type and APPDutch mice.**

A, B. Representative coronal brain sections from APPDutch mice that received a bilateral injection of AAV-VHH-B9 vector at 2 months of age. Brains were harvested 3 weeks post-injection and coronal sections were immunostained for the cMyc tag fused to the VHH. Note the widespread expression of VHH throughout the targeted area.

C, D. Representative coronal brain sections from wild type mice injected with AAV-GFP (C) or APPDutch mice injected with AAV-VHH-B9 (D) and immunostained as indicated for GFP, cMyc or calretinin. GFP and cMyc are pseudocolored green in the merge images. Calretinin is pseudo-colored magenta. Note the diffuse VHH staining pointing to strong intracellular expression. Scale bar, 50 µm.

#### SUPPLEMENTARY METHODS AND MATERIALS

##### Immunization and VHH library construction

Two VHH libraries were generated using a protocol described previously, with minor modifications (Vincke *et al*, 2012). The first library was produced from immunization of a dromedary with recombinant unglycosylated human BACE1 ectodomain (amino acids 46-460) purified from insect cell cultures (Bruinzeel et al., 2002). The second library was produced by immunization of a llama with human BACE1 ectodomain purified from HEK293 cell cultures (Zhou *et al*, 2011). Briefly, the dromedary or llama received six subcutaneous immunizations at weekly intervals with 150 µg BACE1. Immune responses were evaluated using ELISAs, after purification of conventional IgG1 molecules, or the heavy chain-only subclasses IgG2 and IgG3, from serum. In all, 3 different subclasses of IgGs, immunoreactive against BACE1, were detected. The blood of the immunized animal was collected for lymphocyte preparation, followed by RNA isolation and RT-PCR amplification. The cDNAs of the variable fragments of heavy chain-only IgGs were cloned into pHEN4 phagemid and transformed into electrocompetent TG1 *E.coli* cells to generate a library of 6 x $10^7$ transformants with 90% inserts from the dromedary, and another library of 5 x $10^7$ transformants with more than 78% correct inserts from the llama.

##### Panning of VHH libraries

Panning of VHH libraries was performed according to a previously described protocol (Vincke *et al*, 2012), with two different strategies used to enrich BACE1-binding VHHs. The first strategy used an ELISA-based technique combined with high pH elution. Briefly, libraries were rescued with M13K07 helper phage and phage particles were then prepared and applied to ELISA plates pre-coated with purified BACE1 ectodomain. After incubation and washing, the bound phage particles were eluted with 100 mM triethylamine (pH 10.0) and immediately neutralized with 1 M Tris-HCl (pH 7.5). Alternatively, purified BACE1 ectodomain protein was labeled with Sulfo-NHS-SS-Bio (Pierce), according to the manufacturer's protocol. Phage particles were preblocked with 1% (w/w) BSA in panning buffer (50 mM Tris-HCl (pH 7.5), 150 mM NaCl and 0.05% (v/v) Tween-20) and incubated with 200 nM biotin-labeled BACE1. Streptavidin coated paramagnetic beads (Pierce), preblocked in 1% BSA, were used to capture biotin-labeled BACE1. After extensive washes, the bound protein and phage particles were eluted by 50 mM DTT. Eluates from both approaches were used to re-infect exponentially growing *E. coli* TG1 cells for a further round of phage panning. After three consecutive rounds of panning, phages recovered from the second and third rounds were used to infect exponentially growing TG1 cells, which were plated out at dilutions of $10^{-4}$, $10^{-5}$ and $10^{-6}$. Single colonies were picked for further analysis.

**Phage ELISA Screening**

Single colonies retrieved from different rounds of phage panning were grown with shaking at 220 rpm in 2 ml 2x YT medium (1.6% (w/v) bacto tryptone, 0.5% (w/v) NaCl and 1% (w/v) yeast extract in water) supplemented with 100 µg/ml ampicillin and 1% (w/v) glucose in 24-well plates for 8 hours at 37°C. Bacteria in the exponential growth phase were infected using M13K07 helper phages (5 x $10^8$ plaque forming units (pfu) per well) for 20 min at room-temperature. After addition of 70 µg/ml of kanamycin to the cells, they were further grown overnight at 37°C, with constant shaking at 220 rpm. The next morning, bacteria were harvested by centrifugation at 1,200$g_{Av}$ for 20 min. Supernatants containing phage particles were tested for binding to BACE1 in an ELISA. Essentially, 96-well plates (Nunc) were coated with BACE1 ectodomain protein at 100 ng/well overnight at 4°C. Non-coated wells were used as a control. The plates were blocked with 3% milk (w/v) in PBS for 1 hour at RT. After blocking, 100 µl of each phage particle containing supernatant was added to a coated and a non-coated well and incubated for 2 hours at room temperature. Plates were then washed using washing buffer (0.05% (v/v) Tween-20, PBS) and further incubated with HRP-conjugated anti-M13 antibody (GE Healthcare) at a 1:3,000 dilution in 3% (w/v) milk in PBS for 1 hour at room temperature. Plates were then extensively washed and developed using 0.2 mg/ml ABTS (2,2'-azino-bis(3-ethylbenzothiazoline-6-sulfonic acid; Sigma) in 50 mM citric acid, pH 4.0 supplemented with 0.2% (v/v) $H_2O_2$ (Sigma) as a substrate. Signals were read at 405 nm with an ELISA plate reader (2103 EnVision, Perkin Elmer).

**ELISA screening of periplasmic extracts**

The expression vector pHEN4 encodes a PelB (pectate lysase) signal sequence at the N-terminus of the multiple cloning site. Hence, the VHHs encoded by the plasmid are exported to the periplasmic space when expressed in bacterial cultures. To generate periplasmic extract for VHH screening, single colonies, identified by phage ELISA, were inoculated in individual wells of a 24-well plate, each containing 1 ml Terrific Broth (TB) medium (2.4% (w/v) yeast extract, 2% (w/v) tryptone, 0.4% (v/v) glycerol, 17 mM $KH_2PO_4$ and 72 mM $K_2HPO_4$) supplemented with 100 μg/ml ampicillin. Plates were left at 37°C with shaking at 220 rpm. When the $OD_{600}$ reached 0.6 in a well, 1 mM IPTG (isopropyl β-D-1-thiogalactopyranoside) was added to the culture to induce protein expression. Bacteria were grown for approximately 16 hours at 28°C and were then harvested by centrifugation at 1,200$g_{Av}$ for 20 min. Cell pellets were resuspended in TES solution (200 mM Tris-HCl (pH 8), 0.5 mM EDTA, 500 mM sucrose) and incubated on ice for 30 min. To release the VHH from the periplasmic space, a mild osmotic shock was given by adding 1.5x volume of TES buffer diluted 4 times in $H_2O$. After incubation on ice for 45 min, the supernatants were cleared by centrifugation at 1,000$g_{AV}$ for 20 min at 4°C and tested by ELISA. As the VHHs were engineered to contain a C-terminal hemagglutinin (HA) tag, a mouse monoclonal anti-HA antibody (Covance) was used as the detection antibody. An alkaline phosphatase conjugated goat anti-mouse antibody (Sigma) was used as the secondary antibody. ELISA plates were developed using PNPP (p-Nitrophenyl-phosphate; Sigma) as a substrate. Signals were read at 405 nm with an ELISA reader (2103 EnVision, Perkin Elmer).

**Sequence analysis and subcloning of VHHs into a bacterial expression vector**

Positive colonies identified by phage ELISA and periplasmic extract ELISA were analyzed by PCR and restriction enzyme digestion (using HinfI and TaqI enzymes), in order to group different sequence patterns. Primers used for colony PCR were forward 5’-GGACTAGTGCGGCCGCTGGAGACGGTGACCTGGGT-3’ and reverse 5’-TCACACAGGAAACAGCTATGAC-3’. VHH cDNAs with different restriction enzyme digestion patterns were randomly chosen for sequencing and subcloned into the pHEN6 expression vector, which contains the coding sequence for a C-terminal hexa-histidine tag to facilitate purification.

**VHH expression and purification**

pHEN6 expression vectors were transformed into WK6 *E.coli* bacteria. Bacteria were inoculated in 1 litre of TB medium supplemented with 100 μg/ml ampicillin and 0.1% (v/v) glucose. Flasks were incubated at 37°C with continuous agitation at 220 rpm. When the $OD_{600}$ reached 0.75-1.0 in a flask, 1 mM IPTG was added to induce VHH expression. Bacteria were then cultured overnight at 28°C under continuous agitation. The next morning, bacteria were harvested by centrifugation and the periplasmic extract was prepared following the protocol described above. VHHs were purified by affinity chromatography using Ni-NTA beads (Qiagen) and size exclusion chromatography on Superdex-75 (GE Healthcare). Protein concentrations were determined by UV absorption at 280 nm, using the theoretical extinction coefficient of the individual VHHs (calculated based on their amino acid content).

**Affinity measurements**

The binding affinities of selected VHHs for BACE1 were analyzed by surface plasmon resonance (SPR) spectroscopy, using a BIAcore instrument. For immobilization, 2,500 resonance units (RU) purified BACE1 were coupled to a CM5 chip using amine coupling chemistry (EDC/NHS), according to the manufacturer's instructions. Binding/regeneration cycles were performed by injection of the VHHs (in a concentration range of 0-500 nM) at 25°C in HBS buffer (10 mM Hepes (pH 7.5), 150 mM NaCl, 3.5 mM EDTA and 0.005% (v/v) Tween-20) at a constant flow rate of 30 µl/min. Regeneration of the surface was achieved by injection of 10 mM glycine/HCl (pH 1.5). For affinity measurements at acidic pH, running buffer was replaced by a citrate buffer containing 150 mM NaCl, 3.5 mM EDTA, 0.005% (v/v) Tween-20 at pH 4.5.

**Bio-layer interferometry**

Biotinylated human or mouse BACE1 was immobilized at 1 µg/ml for 900 seconds on streptavidin-coated biosensors (Molecular Devices). The biosensors were washed for 120 s in Kinetic Buffer (Molecular Devices) and dipped into wells containing serial dilutions of VHH B9 (1000 s duration). This association step was followed by dipping the biosensors in Kinetic Buffer to allow dissociation (1000 s). All steps were performed at 25°C with constant agitation at 1,000 rpm on an OctetRED96 System (Molecular Devices). Binding parameters were calculated using Molecular Devices Analysis 9.0 software, using a 1:1 homogenous fitting model.

**Production and purification of BACE1:Fc**

The construct expressing human BACE1 (amino acids 1–460) fused to the constant region of human IgG1 (hBACE1:Fc) was described previously (Zhou *et al*, 2011). Mutations including $SQD_{376\text{-}378}$/WAA (F(SQD/WAA)), $EDVATSQDD_{371\text{-}379}$/MGAGLNYE (F(BACE2)), $EVATSQD_{371\text{-}378}$/EGS (ΔF) and $GFPLNQSEVLASVG_{219\text{-}232}$/GAG (ΔA), were introduced into the original construct. Purification of wild type or mutant hBACE1:Fc followed a protocol described previously (Zhou *et al*, 2011). Proteins folded correctly and were active in the MBP-C125APPswe enzymatic assay (see below). Protein purity was assessed by SDS-PAGE using a 4–12% Bis-Tris gel followed by Coomassie staining. Protein concentration was measured by Bradford assay (Bio-Rad). Aliquots of the purified protein were snap-frozen and stored at −80°C.

***In vitro* APP cleavage assay**

The assay was performed essentially as described (Zhou *et al*, 2011), using materials provided by Eli Lilly. Briefly, hBACE1:Fc was diluted to 10 nM in reaction buffer (50 mM ammonium acetate (pH 4.6), 1 mg/ml BSA and 1 mM Triton X-100). The substrate was a fusion protein consisting of maltose binding protein (MBP) and the 125 amino acids from the carboxy terminus of human APP695 containing the Swedish (Sw) mutation (K670M/N671L). Substrate was diluted to 50 nM in reaction buffer. 10 µl test VHH (5 µM) were incubated with 25 µl substrate and 15 µl hBACE1:Fc at 25°C for 3 hours. At the end of the incubation, the reaction mixture was diluted 5-fold in 'stop' buffer (200 mM Tris (pH 8.0), 6 mg/ml BSA and 1 mM Triton X-100). 50 µl of the diluted reaction mixture were then loaded on an ELISA plate pre-coated with an anti-MBP capture antibody. The cleavage product MBP-C26sw was detected by a neo-epitope antibody against the BACE1 cleavage site. Purified MBP-C26sw was used to generate a standard curve. The amount of cleavage product reflects the relative activity of BACE1.

**Dot blot analysis of VHH binding to wild type and mutant BACE1**

200 ng purified wild type or mutant hBACE11-460:Fc (in a final volume of 10 µl) were dotted onto strips of nitrocellulose membrane. After drying, the membrane strips were incubated with blocking buffer (5% w/v milk, 0.05% v/v Tween-20, Tris-buffered saline (TBS: 50 mM Tris-HCl, 150 mM NaCl; pH 7.6)) for 1 h at room temperature. After blocking, the membrane strips were incubated with primary antibodies, including the mouse monoclonal antibodies 1A11, 5G7, 10B8 (Zhou et al., 2011), and VHH-B9 (all diluted to 1 ug/ml in blocking buffer) for 1 h at room temperature. The membranes were then washed three times in washing buffer (0.05% v/v Tween-20, TBS) (5 minutes per wash). After washing, the membrane strip incubated with VHH-B9 was further incubated with an anti-His antibody (1:2000 dilution, Biolegend, J099B12) for 1 h at room temperature, followed by three washes, each of 5 minutes duration. All membranes were subsequently incubated with HRP-conjugated goat-anti-mouse IgG antibody diluted in blocking buffer (1:10,000 dilution, Novus, NB7539) for 1 h at room temperature. After incubation, the membranes were washed 3 times in washing buffer and 2 times in TBS (5 minutes per wash), before signal was developed using enhanced chemiluminescence (ECL) methods. Signals were captured with a Fuji LAS-1000 Luminescent Image Analyzer.

**Cell based assays (cultured neurons and glia)**

Mixed primary brain neurons were derived from C57BL/6J mice at embryonic day 14 (E14), following a previously described protocol (Zhou *et al*, 2011).

Glial cultures were established from C567BL/6J mice at post-natal day 3 (P3), according to a previously described protocol (Voytyuk *et al*, 2018). Once confluent, glia from each dish were frozen in 1 ml freezing medium (90% (v/v) FBS and 10% (v/v) DMSO) to establish a frozen stock. Cultures were reactivated when needed using standard culturing protocols.

*Experiments with neurons using addition of purified nanobodies:*

Generation of recombinant Semliki Forest Viruses (SFV) has been described (Annaert *et al*, 1999). Neurons were cultured in 6 cm dishes (Nunc) and maintained in neurobasal medium (Gibco) supplemented with B27 (Gibco). Neurons were transduced with SFV expressing WT human $APP_{695}$ after three days in culture. Two hours after transduction, neurons were treated with test VHH, diluted in fresh media for 12 hours.

*Experiments with neurons or glia using application of AAV-VHH:*

AAV vectors encoding VHH-B9 or GFP (see below) were used for cell transfection. The pan-BACE inhibitor, Compound J (CpJ), was used as a positive control at a final concentration of 10 nM. DMSO alone was used as the vehicle control. The final amount of vector applied was approximately $3.75 \times 10^{11}$ vector genomes.

AAV vectors were applied to neurons in culture at day 3 *in vitro* (DIV). 3 days post-AAV application, CpJ or DMSO were added according to treatment protocol: i) AAV-VHH-B9 + Vehicle, AAV-GFP + Vehicle (negative control) and AAV-GFP + Compound J (positive control). Media was collected 36 hours later and concentrated using centrifugal concentrators (30 kDa molecular weight cut-off, Millipore). Cells were collected and homogenized in 5 volumes of ice-cold PBS, supplemented with protease (Roche) and phosphatase (Sigma) inhibitors. Lysates were centrifuged in an Ultra Optima TLX (Beckman) at $14{,}000g_{Av}$ and 4°C for 15 min. Supernatants were collected, snap-frozen using liquid $N_2$ and stored at -80°C.

AAV vectors were added to glia when the cells had reached confluency. 3 days post-AAV application, CpJ or DMSO were added as described above. Media and cells were collected 36 hours later and processed as described.

##### *Aβ ELISA*

The concentration of $A\beta_{1-40}$ or $A\beta_{1-42}$ in tissue samples was determined by standard sandwich ELISAs. Monoclonal antibodies $JRFcA\beta_{40/28}$ and $JRFcA\beta_{42/26}$, which recognize the C-terminus of Aβ species terminating at amino acids 40 or 42 respectively, were used for capture. HRP-conjugated JRFAβN/25 antibody, which recognizes the first seven N-terminal amino acids of human Aβ, was used as a detection antibody. Synthetic human $A\beta_{1-40}$ and $A\beta_{1-42}$ peptides were used to generate standard curves. Antibodies were a kind gift from Johnson and Johnson.

##### *Western blot*

The amount of total protein in both culture media and cell extracts was measured using the BCA Method (Thermo Fischer). Samples were prepared by mixing with Laemmli loading buffer and heating at 75°C for 10 min. Samples were then loaded onto 10% Bis-Tris SDS-

PAGE gels (20 µg/lane) and separated by electrophoresis in NuPAGE MES SDS running buffer, except for samples obtained from glial cultures, which were loaded onto 4-12% Bis-Tris SDS-PAGE gels (20 µg/lane) and separated by electrophoresis in NuPAGE MOPS SDS running buffer. Immunoblotting was carried out using standard tank blotting techniques. The membranes were incubated with blocking buffer (5% (w/v) milk, 0.05% (v/v) Tween-20, TBS) for 1 h at room temperature. Primary antibodies in blocking buffer were added overnight at 4°C. Primary antibodies used were WO2 for total Aβ (1:1,000 dilution, The Genetics Company, AB4-10); 6E10 for $sAPP_{\alpha}$ (1:1,000 dilution, Signet Laboratories, #9320-02); anti-$sAPP_{\beta}$ (1:1,000 dilution, Covance, #SIG-39138); custom B63 antibody for full length and carboxy terminal fragments of APP (1:5,000 dilution) (Annaert *et al*, 2001); anti-β-actin (1:3000 dilution, Sigma-Aldrich, A5441), DNER IgG (1:1,000 dilution, R&D Systems, AF2254), anti-llama IgG H&L (1:1,000 dilution, HRP, AB112786), VCAM (1:1,000 dilution, Thermo Fisher Scientific, PA5-47029), SeZ6 (1:1,000, R&D Systems, AF5598) c-Myc (9E10) (1:1,000 dilution, Abcam, ab32), GFP (1:1,000 dilution, Sigma, 11814460001). The membranes were then washed three times in 0.05% (v/v) Tween-20, TBS (7 minutes per wash) and subsequently incubated with corresponding secondary antibody. HRP-conjugated goat-anti-mouse IgG antibody in blocking buffer (1:10,000 dilution, Novus, NB7539), HRP-conjugated goat-anti-rabbit IgG antibody in blocking buffer (1:10,000 dilution, Novus, NBP2-30348H), goat polyclonal anti-mouse IgG-HRP conjugate (1:3,000 dilution, Biorad, 170-6516), goat polyclonal anti-rabbit IgG-HRP conjugate (1:3,000 dilution, Biorad, 170-6515) or rabbit polyclonal anti-goat IgG-HRP conjugate (1:2,000 dilution, DAKO, P0449) for 1 h at room temperature. After the incubation, the membranes were washed 3 times in 0.05% (v/v) Tween-20, TBS and 2 times in TBS (5 minutes per wash) before signal was developed with ECL reagents. Signals were captured and analyzed with a Fuji LAS-1000 Luminescent Image Analyzer.

**AAV vector production**

AAV-based vectors were designed around a standard single stranded AAV expression cassette. The cDNA for the anti-BACE1 VHH-B9 was modified to contain an N-terminal BACE1 signal peptide and a C-terminal cMyc tag. The cDNA sequence encoding VHH-B9 (or GFP for control) was then cloned into the expression cassette, which also contained a cytomegalovirus enhancer/chicken β-actin (CBA) promoter, a woodchuck post-transcriptional regulatory element (WPRE) and a bovine growth hormone poly(A) sequence (pA). AAV-based vectors were then prepared by transfection of HEK293T cells followed by iodixanol purification, as described previously (Fripont *et al*, 2019). HEK293T cells were tri-transfected with an adenovirus helper plasmid, a packaging plasmid encoding an AAV capsid (AAV1 or PHP.B) and the expression cassette. Forty-eight hours after transfection, the cells were harvested and lysed in the presence of sodium deoxycholate (10% (w/v) stock solution) and 50 U/ml benzonase (Sigma) by freeze-thaw cycles. Lysates were cleared by centrifugation and the supernatants containing released vector were recovered. Vectors were subsequently purified using discontinuous iodixanol gradients (Sigma) and HiTrap HQ columns (GE Healthcare). The genomic titer of each vector was determined by quantitative PCR and expressed as vector genomes (vg) per ml. AAV-PHP.B-based vectors were used in tissue culture experiments. AAV1-based vectors were used for *in vivo* administration.

**Animals**

All animal procedures were performed in accordance with the regulations of the Institutional Animal Care and Use Committee of KU Leuven. Animals were kept in a specific pathogen free (SPF) facility with controlled humidity, temperature and light conditions. Food and water were provided *ad libitum*.

**Stereotaxic injections, sample preparation and analysis**

APPDutch mice (Herzig *et al.*, 2004) used in the treated and control groups received a direct intraparenchymal delivery of AAV vector (bilateral injection of 2 x $10^{10}$ vg per injection site) using a 30-gauge needle (Hamilton Neuro Syringes, model 7002 KH). The stereotaxic coordinates (from bregma) were A/P: -2.4, M/L: +/-2.6, V/L: -2.5. Animals were euthanized 3 weeks post-injection.

For biochemical analyses, animals were perfused with ice-cold PBS before hippocampi were dissected out from the brain and flash-frozen in liquid $N_2$. Samples were stored at -80°C until use. Soluble proteins were extracted on ice from thawed samples, using mechanical homogenization in lysis buffer (TBS supplemented with complete protease inhibitor, Roche). Briefly, 30 mg of tissue was added to 300 µl lysis buffer and disrupted using 10 strokes of a motor-driven glass-Teflon homogenizer at 800 rpm. Samples were then transferred to 1.5 ml microtubes (Sarstedt) and cleared by centrifugation at 17,000$g_{Av}$ for 30 min at 4°C. The supernatant (soluble fraction) was collected, snap-frozen using liquid $N_2$ and stored at -80°C until further use. A fraction of the supernatant was later diluted and analyzed for Aβ levels by ELISA (see above).

For histological analyses, animals were transcardially perfused with PBS, followed by 4% paraformaldehyde (PFA) in PBS. Brains were collected and post-fixed for a further 24 hours in 4% PFA. Vibratome sections of 50 µm thickness were cut using a Leica VT1000S. Free-floating vibratome sections were blocked for 1 to 2 hours at room temperature in 1% (v/v) Triton X-100 in PBS, supplemented with 10% normal donkey serum (NDS). For BACE1 staining, antigen retrieval was performed, prior to blocking, by boiling sections in citrate buffer (10 mM citric acid, 0.05% (v/v) Tween-20; pH 6.0 NaOH) followed by three washes in PBS, each of five minutes duration. Sections were then incubated overnight at 4°C in primary antibodies diluted in blocking solution. Primary antibodies used for staining were rabbit anti-GFP (1:400 dilution, Synaptic Systems, #132002), rat anti-cMyc (1:400 dilution, Bio-Rad, #MCA1929), guinea pig anti-calretinin (Calbindin D29k) (1:1,000 dilution, Synaptic Systems, #214104), rabbit anti-BACE1 (1:150 dilution, Cell Signaling, D10E5). Sections were then washed three times in PBS, with each wash lasting for 10 minutes. Sections were then incubated for 2 hours at room temperature with secondary antibodies diluted in 1% (v/v) Triton-X100 in PBS. Secondary antibodies used were donkey anti-rabbit Alexa 488 (1:1,000 dilution, Invitrogen, #R37118), donkey anti-rabbit Cy3 (1:200 dilution, Jackson Immuno, #711-165-152), donkey anti-rat Alexa 488 (1:200 dilution, Jackson Immuno, #712-545-150) and donkey anti-guinea pig Cy3 (1:200 dilution, Jackson Immuno, #712-095-150). Sections were then given three washes in PBS, with each wash lasting for 10 minutes. Slices were mounted onto microscopy slides, using Fluoromount G containing DAPI (SouthernBiotech). Wide field images were taken using a Zeiss Slide scanner (Axio Scan Z1) with a PL APO 20x/NA 0.8 objective, controlled using Zen 2 data acquisition software (Appendix Figure S4A, B). Other images were taken on a Leica DM5500 epifluorescence microscope (Leica Microsystems, Wetzlar, Germany) with a PL APO 10x/NA 0.4 objective (Appendix Figure S4C, D). Confocal images were taken using a Leica SP8 microscope with a HC PL APO CS2 40x/1.30 objective (Figure 3A-D).

**REFERENCES**

Annaert WG, Esselens C, Baert V, Boeve C, Snellings G, Cupers P, Craessaerts K & De Strooper B (2001) Interaction with telencephalin and the amyloid precursor protein predicts a ring structure for presenilins. *Neuron* **32:** 579–589

Annaert WG, Levesque L, Craessaerts K, Dierinck I, Snellings G, Westaway D, George-Hyslop PS, Cordell B, Fraser P & De Strooper B (1999) Presenilin 1 controls gamma-secretase processing of amyloid precursor protein in pre-golgi compartments of hippocampal neurons. *J. Cell Biol.* **147:** 277–294

Fripont S, Marneffe C, Marino M, Rincon MY & Holt MG (2019) Production, Purification, and Quality Control for Adeno-associated Virus-based Vectors. *J Vis Exp*

Vincke C, Gutiérrez C, Wernery U, Devoogdt N, Hassanzadeh-Ghassabeh G & Muyldermans S (2012) Generation of single domain antibody fragments derived from camelids and generation of manifold constructs. *Methods Mol. Biol.* **907:** 145–176

Voytyuk I, Mueller SA, Herber J, Snellinx A, Moechars D, van Loo G, Lichtenthaler SF & De Strooper B (2018) BACE2 distribution in major brain cell types and identification of novel substrates. *Life Sci Alliance* **1:** e201800026

Zhou L, Chávez-Gutiérrez L, Bockstael K, Sannerud R, Annaert W, May PC, Karran E & De Strooper B (2011) Inhibition of beta-secretase in vivo via antibody binding to unique loops (D and F) of BACE1. *J. Biol. Chem.* **286:** 8677–8687
