## Appendix Figure S1 for "AAV mediated delivery of a novel anti-BACE1 VHH reduces Abeta in an Alzheimer’s disease mouse model"

**Appendix Figure S1. Rincon MY., Zhou L., *et al.***

```
            <  --  Framework-1  -- >  < CDR1 > < --Framework-2 >
VHH-B9      DVQLQESGGGSVQAGGSLRLSCAAS EYTYGYCS MGWYRQAPGKERELVST 50
VHH-10C4    QVQLQESGGGSVQAGGFLRLSCAAS GYTYSTCS MAWYRQAPGKERELVSS 50
VHH-4A2     QVQLQESGGGLVQPGGSLRLSCAAS GFTFETQY MTWVRQAPGKGPEYVSS 50

            < CDR2> < ----------- Framework-3 -----------   >
VHH-B9      ITSDGST S-YVD-SVKGRFTISQDNAKNTVYLQMNSLKPEDTAKYYC 98
VHH-10C4    IRNDGST A-YAD-SVKGRFTISQDNAKNTVYLQMNSLKPEDTAMYYC 98
VHH-4A2     INSGGTI KYYANSSVKGRFTISRDNAKNTLYLQMNNLRPEDTAIYYC 100

            <  ---- CDR3 ----  > <Framework-4>
VHH-B9      YTK-----TCANKLGAKFIS WGQGTQVTVSS 121
VHH-10C4    NIR-IGVGP-GGTCSIYAPY WGEGTQVTVSS 124
VHH-4A2     QLGQWAG-------VGAASS RGQGTQVTVSS 121
```
