## Supplementary figures and images for "AAV mediated delivery of a novel anti-BACE1 VHH reduces Abeta in an Alzheimer’s disease mouse model"

### Appendix Figure S2

Appendix Figure S2. Rincon MY., Zhou L., *et al.*

A

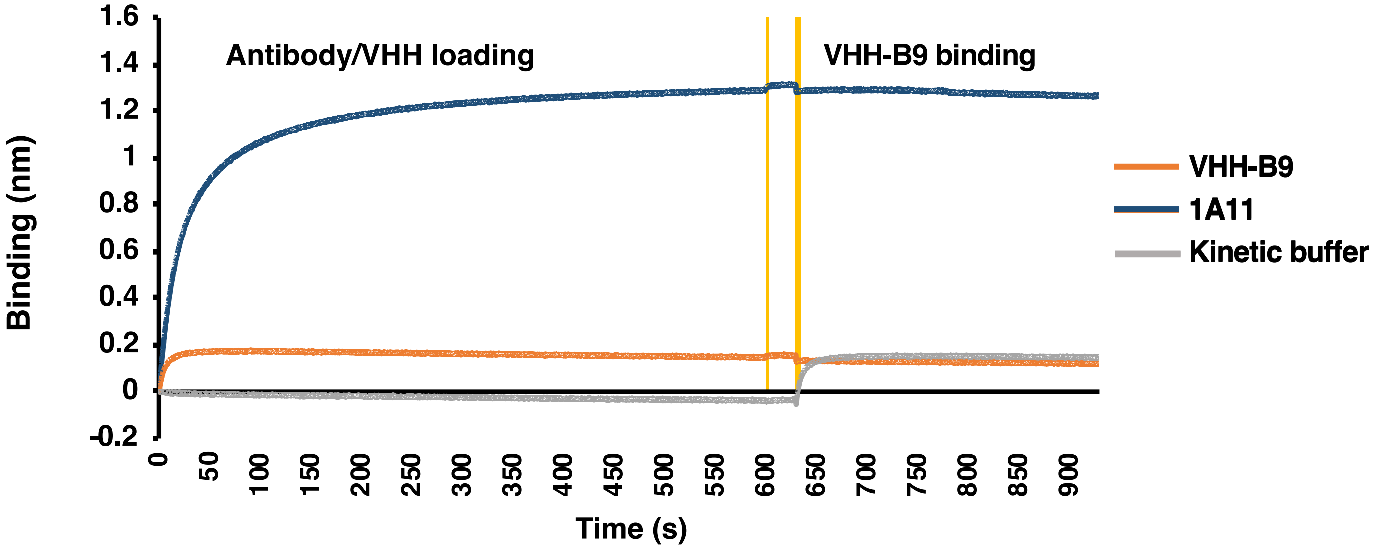


B

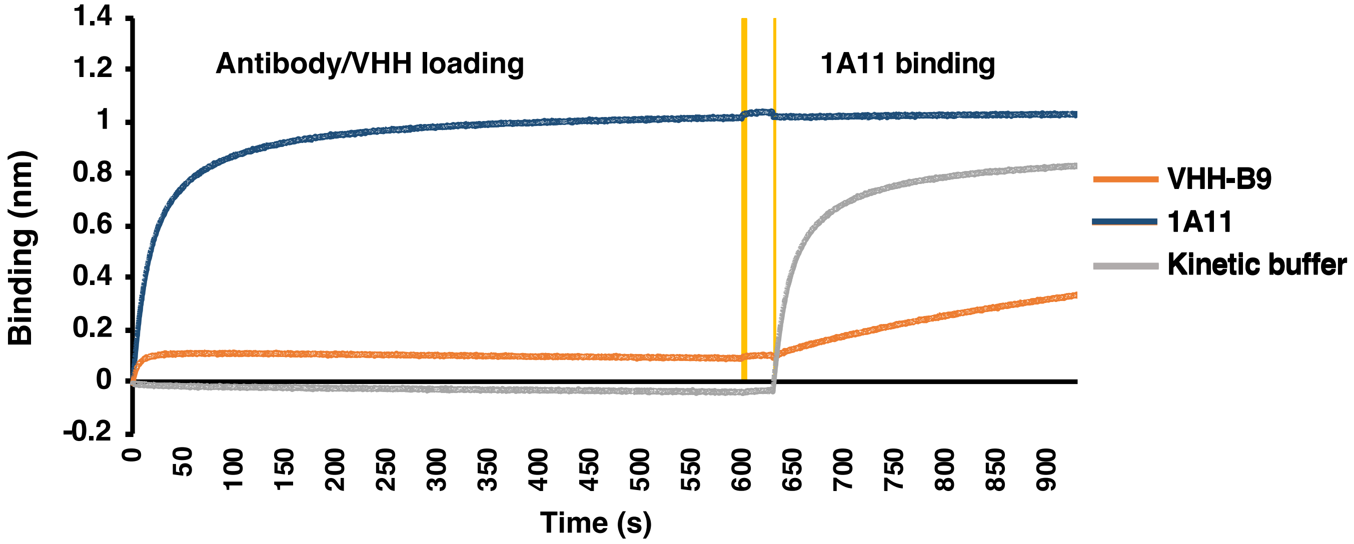

### Appendix Figure S3

**Appendix Figure S3. Rincon MY., Zhou L., *et al.***

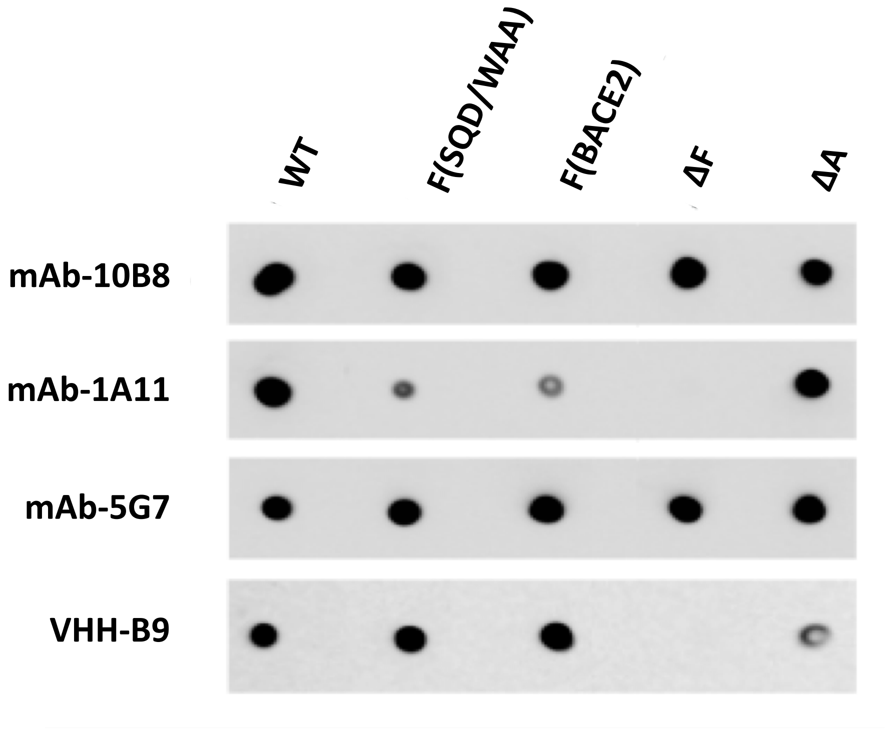

### Appendix Figure S4

**Appendix Figure S4. Rincon MY., Zhou L., *et al.***

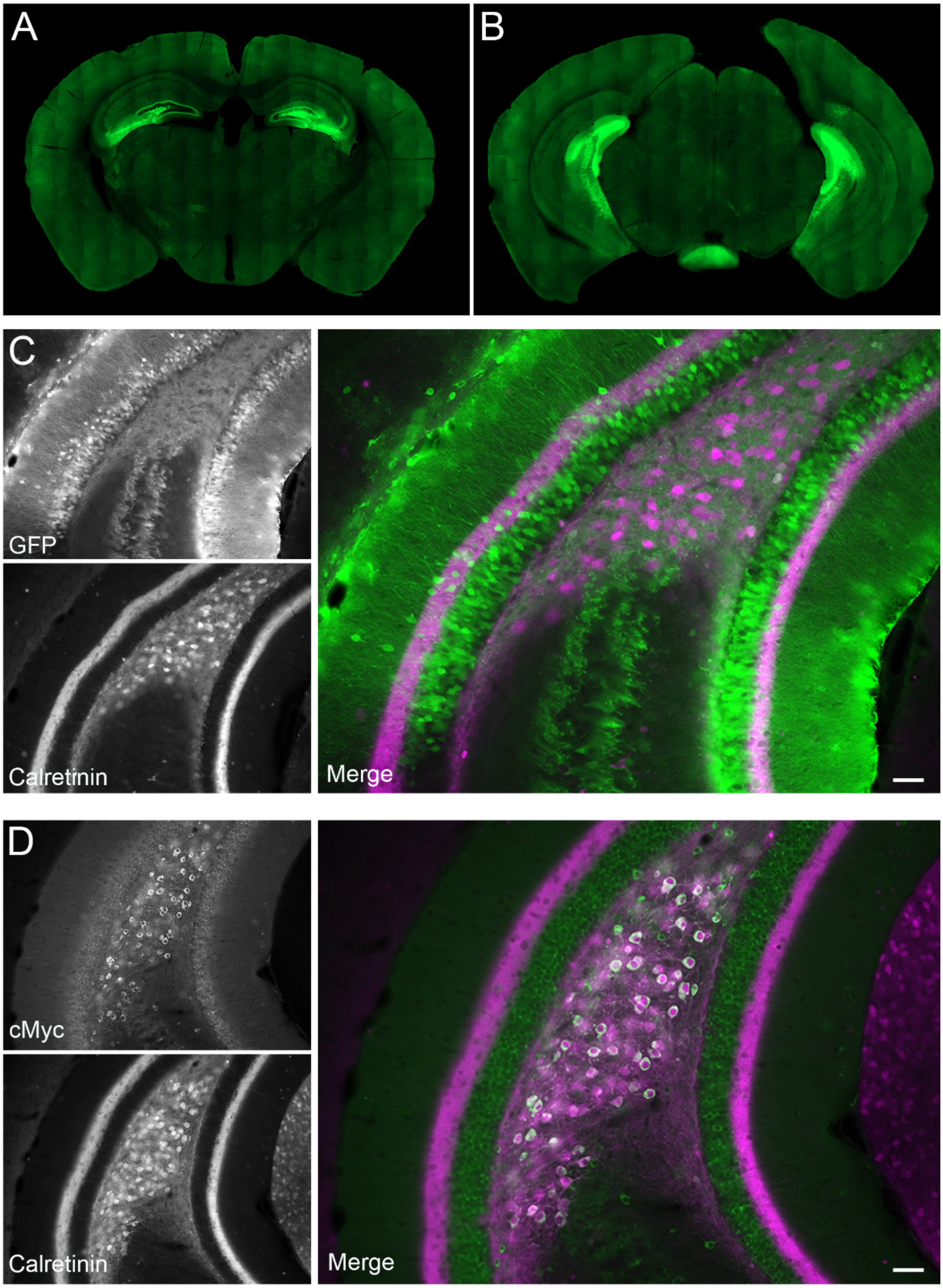
